# Multi-Region Neural Representation: A novel model for decoding visual stimuli in human brains

**DOI:** 10.1101/097675

**Authors:** Muhammad Yousefnezhad, Daoqiang Zhang

## Abstract

Multivariate Pattern (MVP) classification holds enormous potential for decoding visual stimuli in the human brain by employing task-based fMRI data sets. There is a wide range of challenges in the MVP techniques, i.e. decreasing noise and sparsity, defining effective regions of interest (ROIs), visualizing results, and the cost of brain studies. In overcoming these challenges, this paper proposes a novel model of neural representation, which can automatically detect the active regions for each visual stimulus and then utilize these anatomical regions for visualizing and analyzing the functional activities. Therefore, this model provides an opportunity for neuroscientists to ask this question: what is the effect of a stimulus on each of the detected regions instead of just study the fluctuation of voxels in the manually selected ROIs. Moreover, our method introduces analyzing snapshots of brain image for decreasing sparsity rather than using the whole of fMRI time series. Further, a new Gaussian smoothing method is proposed for removing noise of voxels in the level of ROIs. The proposed method enables us to combine different fMRI data sets for reducing the cost of brain studies. Experimental studies on 4 visual categories (words, consonants, objects and nonsense photos) confirm that the proposed method achieves superior performance to state-of-the-art methods.

## 1 Introduction

A universal unanswered question in neuroscience is how the human brain activities can be mapped to the different brain tasks? As one of the main techniques in task-based functional Magnetic Resonance Imaging (fMRI) analysis, Multivariate Pattern (MVP) is a conjunction between neuroscience and computer science, which can extract and decode brain patterns by applying the classification methods [1, 2]. Indeed, it can predict patterns of neural activities associated with different cognitive states [3, 4] and also can define decision surfaces to distinguish different stimuli for decoding the brain and understanding how it works [5, 6]. Analyzing the patterns of visual objects is one of the most interesting topics in MVP classification, which can enable us to understand how brain stores and processes the visual stimuli. It can be used to find novel treatments for mental diseases or even to create a new generation of the user interface.

Technically, MVP classification is really a challenging problem. Firstly, most of the fMRI data sets are noisy and sparse, which can decrease the performance of MVP methods [7]. The next challenge is defining the regions of interest (ROIs) [4]. As mentioned before, fMRI techniques allow us to study what information are represented in the different regions. So, it is really important to know what are the effects of different stimuli on the brain regions, especially in complex tasks (doing some simple tasks at the same time such as watching photos and tapping keys). On the one hand, most of the previous studies manually selected the ROIs. On the other hand, defining wrong ROIs can significantly decrease the performance of MVP methods [3, 4]. Another challenge is the cost of brain studies. Combining different homogeneous fMRI data sets can be considered as a solution for this problem but data must be normalized in a standard space. The procedure of normalization can increase the time and space complexities and decrease the robustness of MVP techniques, especially in voxel-based methods [5]. The last challenge is visualization. As a machine learning technique, MVP represents the numerical results in the voxel-level, network connections, etc. Sometimes, it is so hard for neuroscientists to find a relation between the generated results and the cognitive states.

The contributions of the paper are four fold: firstly, the proposed method estimates and analyzes a snapshot of brain image for each stimulus based on the level of using oxygen in the brain instead of analyzing whole of fMRI time series. Indeed, employing these snapshots can dramatically decrease the sparsity. Secondly, our methods can automatically detect active regions for each stimulus and dynamically define ROIs for each data set. Further, it develops a novel model of neural representation for analyzing and visualizing functional activities in the form of anatomical regions. This model can provide a compact and informative representation of neural activities for neuroscientists to understand: what is the effect of a stimulus on each of the automatically detected regions instead of just study the fluctuation of a group of voxels in the manually selected ROIs. The next contribution is a new Gaussian smoothing method for removing noise of voxels in the level of anatomical regions. Lastly, this paper employs the L1-regularization Support Vector Machine (SVM) [8] method for creating binary classification at the ROIs level and then combine these classifiers by using the Bagging algorithm [9, 10] for generating the MVP model.

## 2 Related Works

As the most prevalent techniques in the human brain decoding, MVP methods can predict patterns of neural activities. Since spatial resolution and within-area patterns of response in fMRI can provide an informative representation of stimulus distinctions, most of previous MVP studies for decoding the human brain focused on task-based fMRI data sets [5]. They used these data sets for generating different forms of neural representation, include usually voxels (volume elements in brain images), nodes on the cortical surface, the average signal for an area, a principal or independent component, or a measure of functional connectivity between a pair of locations [5, 6, 11, 12]. Previous studies demonstrated that MVP classification can also distinguish many other brain states such as recognizing visual [5, 6, 11], or auditory stimuli [13].

Pioneer studies just focused on the special regions of the human brain, such as the Fusiform Face Area (FFA) or Parahippocampal Place Area (PPA) [11]. Haxby et al. showed that different visual stimuli, i.e. human faces, animals, etc., represent different responses in the brain [5, 14]. Hanson et al. developed combinatorial codes in the ventral temporal lobe for object recognition [6]. Norman et al. argued for using SVM and Gaussian Naive Bayes classifiers [15]. Anderson and Oates studied the chance of applying non-linear Artificial Neural Network (ANN) on brain responses [1].

There is great potential for employing sparse methods for brain decoding problems [16, 17]. Carroll et al. employed the Elastic Net [18] for prediction and interpretation of distributed neural activity with sparse models [19]. Richiardi et al. extracted the characteristic connectivity signatures of different brain states to perform classification [20]. Varoquaux et al. proposed a small-sample brain mapping by using sparse recovery on spatially correlated designs with randomization and clustering. Their method is applied on small sets of brain patterns for distinguishing different categories based on a one-versus-one strategy [21]. McMenamin et al. studied subsystems underlie abstract-category (AC) recognition and priming of objects (e.g., cat, piano) and specific-exemplar (SE) recognition and priming of objects (e.g., a calico cat, a different calico cat, a grand piano, etc.). Technically, they applied SVM on manually selected ROIs in the human brain for generating the visual stimuli predictors [4]. Mohr et al. compared four different classification methods, i.e. L1/2 regularized SVM [8, 22], the Elastic Net, and the Graph Net [23], for predicting different responses in the human brain. They show that L1-regularization can improve classification performance while simultaneously providing highly specific and interpretable discriminative activation patterns [3]. Osher et al. proposed a network (graph) based approach by using anatomical regions of the human brain for representing and classifying the different visual stimuli responses (faces, objects, bodies, scenes) [12].

## 3 The Proposed Method

The fMRI techniques visualize the neural activities by measuring the level of oxygenation or deoxygenation in the human brain, which is called Blood Oxygen Level Dependent (BOLD) signals. Technically, these signals can be represented as time series for each subject. Most of the MVP techniques directly analyze these noisy and sparse time series for understanding which patterns are demonstrated for different stimuli.

The main idea of our proposed method is so simple. Instead of analyzing whole of the time series, the proposed method estimates and analyzes a snapshot of brain image for each stimulus when the level of using oxygen is maximized. As a result, this method can automatically decrease the sparsity of brain image. The proposed method is applied in three stages: firstly, snapshots of brain image are selected by finding local maximums in the smoothed version of the design matrix. Then, features are generated in three steps, including normalizing to standard space, segmenting the snapshots in the form of automatically detected anatomical regions, and removing noise by Gaussian smoothing in the level of ROIs. Finally, decision surfaces [5] are generated by utilizing the bagging method on binary classifiers, which are created by applying L1-regularized SVM on each of neural activities in the level of ROIs.

### 3.1 Snapshots Selection

fMRI time series collected from a subject can be denoted by **F** ∈ ℝ^*t*×*m*^, where *t* is the number of time samples, and *m* denotes the number of voxels. Same as previous studies [1, 4, 3, 11], **F** can be formulated by a linear model as follows:

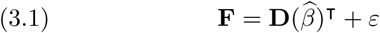

where **D** ∈ ℝ^*t*×*p*^ denotes the design matrix, *ε* is the noise (error of estimation), 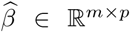 denotes the sets of correlations (estimated regressors) between voxels. The design matrix can be denoted by **D** = {**d**_1_,**d**_2_,…,**d**_i_,…, **d**_p_}, and the sets of correlations can be defined by 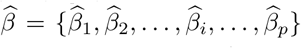. Here, **d***_i_* ∈ ℝ*^t^* and 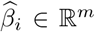 are the column of design matrix and the set of correlations for *i* – *th* category, respectively. *p* is also the number of all categories in the experiment **F**. In fact, each category (independent tasks) contains a set of homogeneous visual stimuli. In addition, the nonzero voxels in 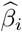 represents the location of all active voxels for the *i*–*th* category [24]. As an example, imagine during a unique session for recognizing visual stimuli, if a subject watches 4 photos of cats and 3 photos of houses, then the design matrix contains two columns; and there are also two sets of correlations between voxels, i.e. one for watching cats and another for watching houses. Indeed, the final goal of this section is extracting 7 snapshots of the brain image for the 7 stimuli in this example.

##### Algorithm 1

The Snapshots Selection Algorithm

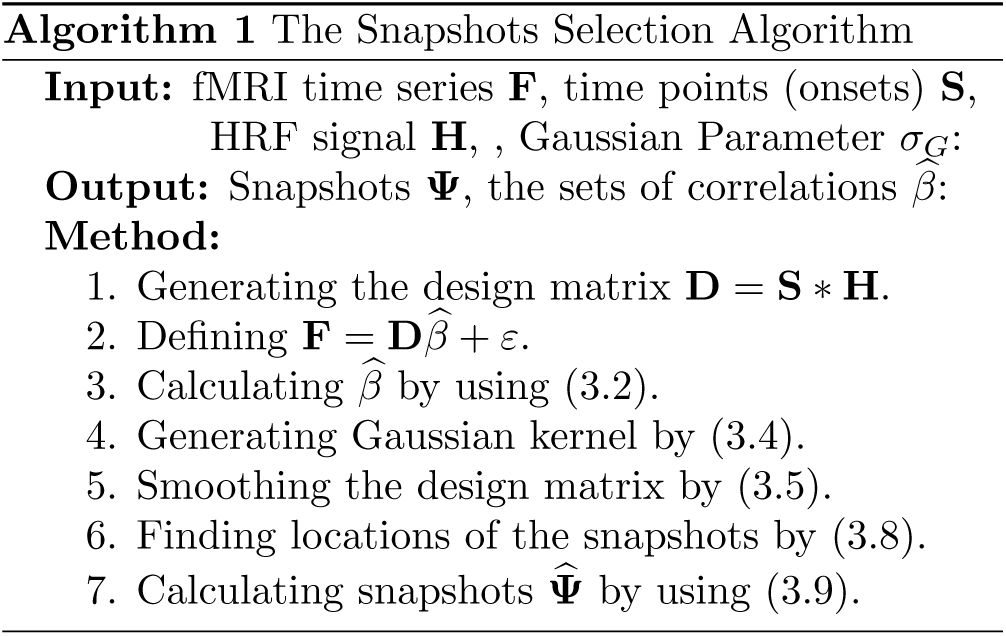

The design matrix can be classically calculated by convolution of time samples (or onsets: **S** = { **S**_1_, **S**_2_,…,**S**_i_,…,**S**_p_}) and **H** as the Hemodynamic Response Function (HRF) signal, **d**_i_ = **S**_i_ \***H** ⇒ **D** = **S** \***H** [11, 24]. In addition, there is a wide range of solutions for estimating 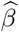 values. This paper uses the classical method Generalized Least Squares (GLS) [24] for estimating the 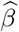 values where ∑ is the covariance matrix of the noise (*V ar*(*ε*) = ∑σ^2^ ≠ 𝕀 σ_2_):

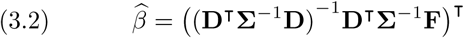

Each local maximum in **d**_*i*_ represents a location where the level of using oxygen is so high. In other words, the stimulus happens in that location. Since **d**_*i*_ mostly contains small spikes (especially for event-related experiments), it cannot be directly used for finding these local maximums. Therefore, this paper employs a Gaussian kernel for smoothing the **d**_*i*_ signal. Now, the interval 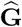 is defined as follows for generating the kernel:

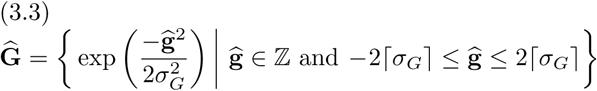

where *σ_G_* > 0 denotes a positive real number; ⌈·⌉ is the ceiling function; and ℤ denotes the set of integer numbers. Gaussian kernel is also defined by normalizing 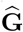 as follows:

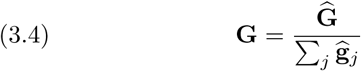

where 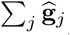 is the sum of all elements in the interval 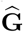. This paper defines the smoothed version of the design matrix by applying the convolution of the Gaussian kernel **G** and each column of the design matrix (**d**_i_) as follows:

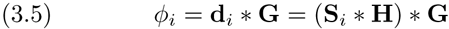

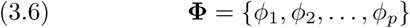

where *ϕ*_i_ = *f* (**S**_*i*_, **H**, **G**). Since the level of smoothness in **Φ** is related to the positive value in (3.3), *σ_G_* = 1 is heuristically defined to generate the optimum level of smoothness in the design matrix. The general assumption here is the 0 < *σ_G_* < 1 can create design matrix, which is sensitive to small spikes. Further, *σ_G_* > 1 can rapidly increase the level of smoothness, and remove some weak local maximums, especially in the event-related fMRI data sets. Figure 1 illustrates two examples of the smoothed columns in the design matrix. The local maximum points in the *ϕ_i_*, can be calculated as follows:

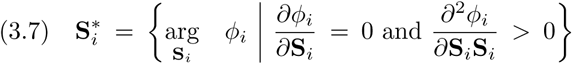

where 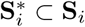 denotes the set of time points for all local maximums in *ϕ_i_*. The sets of maximum points for all categories can be denoted as follows:

**Figure 1:**
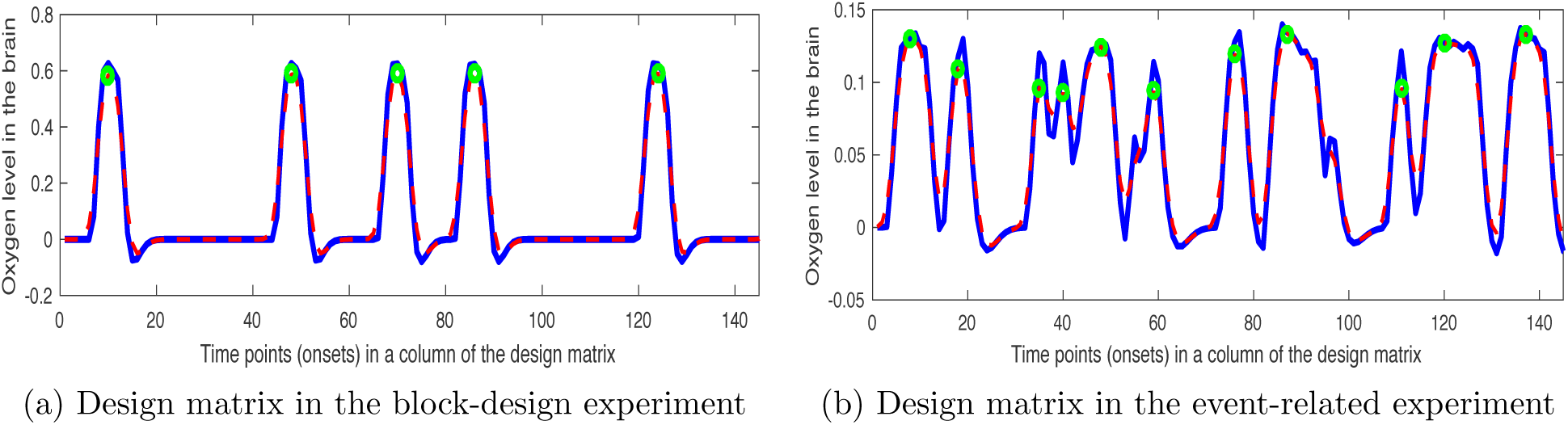
Two examples of smoothed version of the design matrix. The blue lines show the original convolution (**d**_*i*_ = **S**_*i*_ * **H**), the red dashed lines depict the smooth versions (*ϕ_i_* = (**S**_*i*_ * **H**) * **G**), and the green circles illustrate the locations 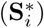 of the detected snapshots 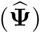.

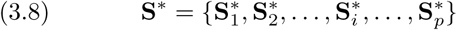

As mentioned before, the fMRI time series can be also denoted by 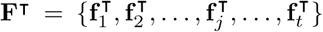, where 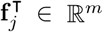 is all voxels of fMRI data set in the *j* – *th* time point. Now, the set of snapshots can be formulated as follows:

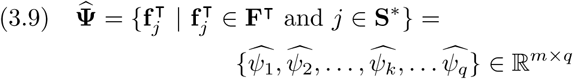

where *q* is the number of snapshots in the brain image **F**, and 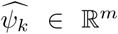 denotes the snapshot for *k* – *th* stimulus. These selected snapshots are employed in next section for extracting features of the neural activities. Algorithm 1 illustrates the whole of procedure for generating the snapshots from the time series **F**.

### 3.2 Feature Extraction

In this paper, the feature extraction is applied in three steps, i.e. normalizing snapshots to standard space, segmenting the snapshots in the form of automatically detected regions, and removing noise by Gaussian smoothing in the level of ROIs. As mentioned before, normalizing brain image to the standard space can increase the time and space complexities and decrease the robustness of MVP techniques, especially in voxel-based methods [5]. On the one hand, most of the previous studies [3, 4, 6, 11] preferred to use original data sets instead of the standard version because of the mentioned problem. On the other hand, this mapping can provide a normalized view for combing homogeneous data sets. As a result, it can significantly reduce the cost of brain studies and rapidly increase the chance of understanding how the brain works. Employing brain snapshots rather than analyzing whole of data can solve the normalization problem.

Normalization can be formulated as a mapping problem. Indeed, brain snapshots are mapped from ℝ^*m*^ space to the standard space ℝ^*n*^ by using a transformation matrix for each snapshot. There is also another trick for improving the performance of this procedure. Since the set 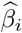 denotes the locations of all active voxels for the *i* – *th* category, it represents the brain mask for that category and can be used for generating the transform matrix related to all snapshots belong to that category. For instance, in the example of the previous section, instead of calculating 7 transform matrices for 7 stimuli, we calculate 2 matrices, including one for the category of cats and the second one for the category of houses. This mapping can be denoted as follows:

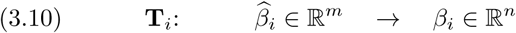

where **T**_*i*_ ∈ ℝ^*m*×*n*^ denotes the transform matrix, 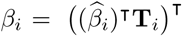 is the set of correlations in the standard space for *i* – *th* category. This paper utilizes the FLIRT algorithm [25] for calculating the transform matrix, which minimizes the following objective function:

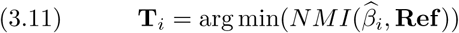

where the function *NMI* denotes the Normalized Mutual Information between two images [25], and **Ref** ∈ ℝ^*n*^ is the reference image in the standard space. This image must contain the structures of the human brain, i.e. white matter, gray matter, and CSF. These structures can improve the performance of mapping between the brain mask in the selected snapshot and the general form of a standard brain. The performance of (3.11) will be analyzed in the supplementary materials^1^. In addition, the sets of correlations for all of categories in the standard space is denoted by *β* = {*β*_1_, *β*_2_,…, *β_i_*,…, *β_p_*} ∈ ℝ^*n*×*p*^, and the sets of transform matrices is defined by **T** = {**T**_1_, **T**_2_,…, **T**_*i*_,…, **T***_p_*}. Now, the *Select* function is denoted as follows to find suitable transform matrix for each snapshot:

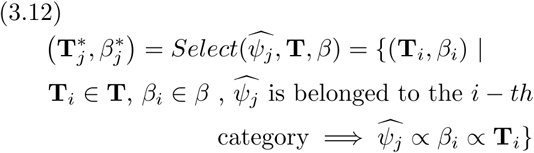

where 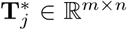 and 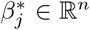 are the transform matrix and the set of correlations related to the *j* – *th* snapshot, respectively. Based on (3.12), each normalized snapshot in the standard space is defined as follows:

##### Algorithm 2

The Feature Extraction Algorithm

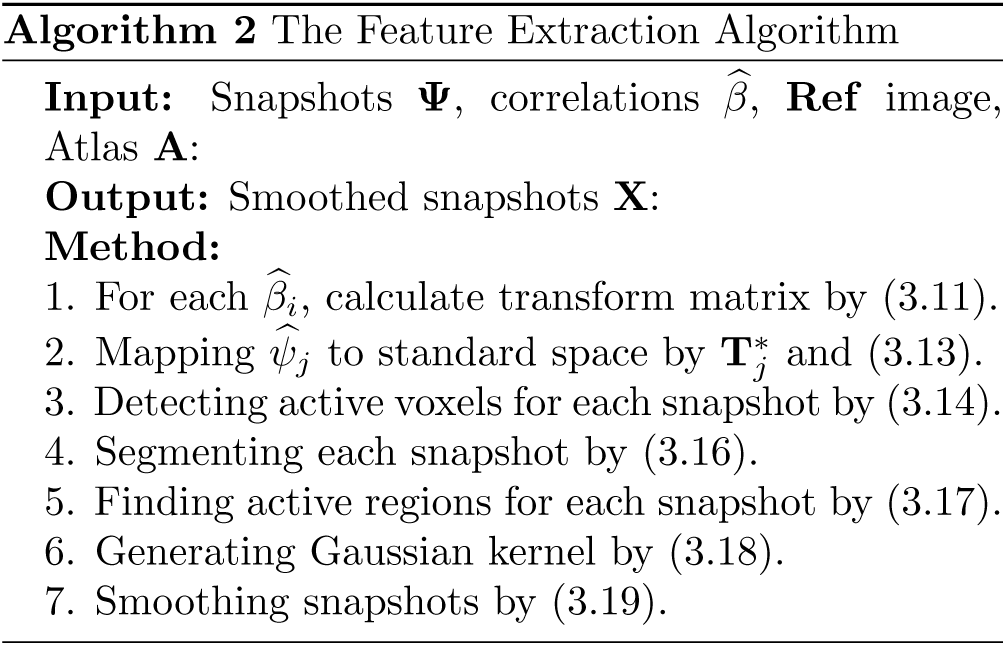

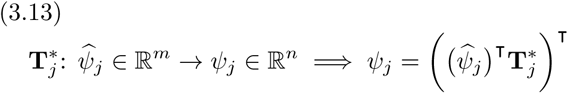

where *ψ_j_* ∈ ℝ^*n*^ is the *j* – *th* snapshot in the standard space. Further, all snapshots in the standard space can be defined by **Ψ** = {*ψ*_1_, *ψ*_2_,…, *ψ_j_*,…, *ψ_q_*} ∈ ℝ^*n*×*q*^. As mentioned before, nonzero values in the correlation sets depict the location of the active voxels. Based on (3.12), this paper uses these correlation sets as weights for each snapshot as follows:

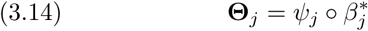

where ○ denotes Hadamard product, and **Θ**_*j*_ ∈ ℝ^*n*^ is the *j* – *th* modified snapshot, where the values of deactivated voxels (and also deactivated anatomical regions) are zero in this snapshot. As the final product of normalization procedure, the set of snapshots can be denoted by **Θ** = {**Θ**_1_, **Θ**_2_,…,**Θ**_*j*_,…,**Θ**_*q*_}. Further,each snapshot can be defined in the voxel level as follows, where 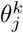 is the *k* – *th* voxel of *j* – *th* snapshot:

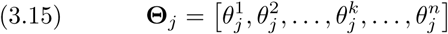

The next step is segmenting the snapshots in the form of automatically detected regions. Now, consider anatomical atlas **A** ∈ ℝ^*n*^ = {**A**_1_, **A**_2_,…, **A**_*l*_,…, **A**_*L*_}, where 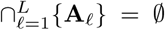, 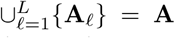, and *L* is the number of all regions in the anatomical atlas. Here, **A**_*ℓ*_ denotes the set of voxel locations in the snapshots for the *ℓ* – *th* anatomical region. A segmented snapshot based on the *ℓ* – *th* region can be denoted as follows:

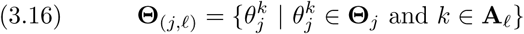

where **Θ**_(*j*,*ℓ*)_ ⊂ **Θ**_*j*_ is the subset of voxels in the snapshot **Θ**_*j*_, which these voxels are belonged to the the *ℓ* – *th* anatomical region. In addition, the sets of all anatomical regions in the *j* – *th* snapshot can be defined by **Θ**_*j*_ = { **Θ**(_*j*,1_) ∪ **Θ**(_*j*,2_) ∪ … ∪ **Θ**(_*j*,*ℓ*_) ∪ … ∪ 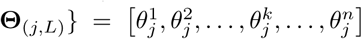. The automatically detected active regions can be alsodefined as follows:

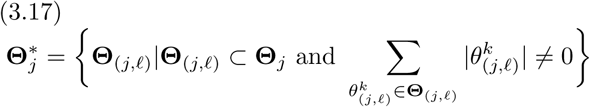

where 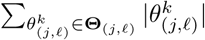 represents sum of all voxels in the **Θ**(_*j*,*ℓ*_). Based on (3.17), active regions in the *j* – *th* snapshot can be defined as the regions with non-zero voxels because values of all deactivated voxels are changed to zero by using (3.14). The last step is removing noise by Gaussian smoothing in the level of ROIs. As the first step, a Gaussian kernel for each anatomical region can be defined as follows:

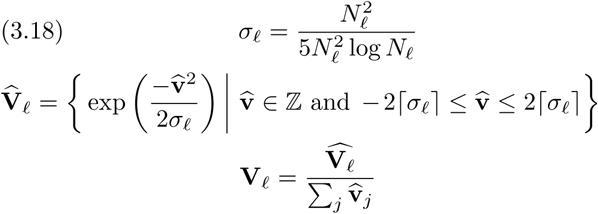

where *N_ℓ_* denotes the number of voxels in *ℓ* – *th* region, and 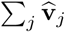 is sum of all values in the interval 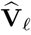. Indeed, the level of smoothness is related to *σ_ℓ_*, which is heuristically calculated for each region based on the number of voxels in that region. As the second step, the smoothed version of the *j* – *th* snapshot can be defined as follows:

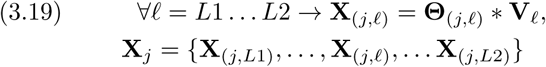

where 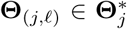 is the *ℓ* – *th* active region of *j* – *th* snapshot, and * denotes the convolution between the active region and the Gaussian kernel related to that region. Further, *L*1 and *L*2 are the first and the last active regions in the snapshot, where 1 ≤ *L*1 ≤ *L*2 ≤ *L*. Figure 2 demonstrates two examples of smoothed anatomical regions in the voxel level. All smoothed snapshots can be defined by **X** = {**X**_1_, **X**_2_,…, **X**_*j*_,…, **X**_*q*_}. Moreover, Algorithm 2 shows the whole of procedure for extracting features.

**Figure 2:**
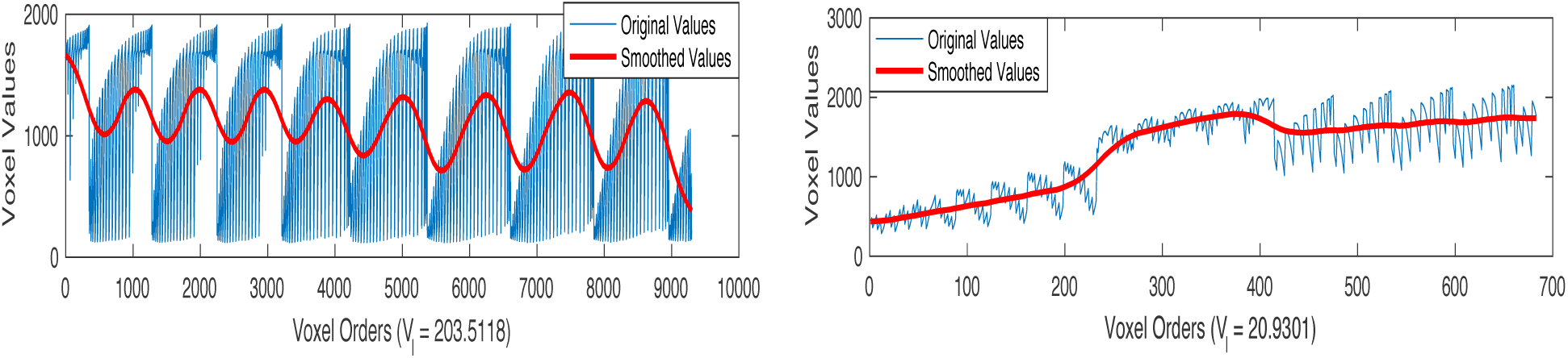
Two examples of smoothed anatomical regions (**X**_(*j*,*ℓ*)_) in the voxel level. Blue lines are the original data, and red lines depict the smoothed values.

### 3.3 Classification Method

As a classical classification method, Support Vector Machine (SVM) [8, 22] decreases the operating risk and can find an optimized solution by maximizing the margin of error. As a result, it can mostly generate better performance in comparison with other methods, especially for binary classification problems. Therefore, SVM is generally used in the wide range of studies for creating predictive models [4, 3, 6, 11]. The final goal of this section is employing the L1-regularization SVM [8] method for creating binary classification at the ROIs level, and then combining these classifiers by using the Bagging algorithm [9, 10] for generating the MVP final predictive model.

As mentioned before, fMRI time series for a subject can be denoted by **F**. Since fMRI experiment is mostly multi-subject, this paper denotes **F**_u_, = 1:*U* as fMRI time series (sessions) for all subjects, where *U* is the number of subjects. In addition, 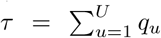 is defined as the number of all snapshots in a unique fMRI experiment. Here *q_u_* is the number of snapshots for *u* – *th* subject. Further, the original ground truth (the title of stimuli such that cats, houses, etc.) for all snapshots is denoted by **Y** = {*y*_1_, *y*_2_,…,*y_j_*,…*y_τ_*}, where *y_j_* denotes the ground truth for *j* – *th* snapshot. Since this paper uses a one-versus-all strategy, we can consider that *y*_j_ ∈ {–1, +1}. This paper applies following objective function on automatically detected active regions as the L1-regularization SVM method for creating binary classification in the level of ROIs [3, 8]:

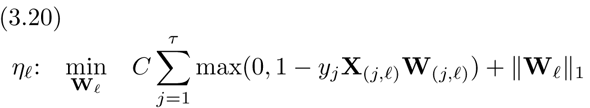

where *C* > 0 is a real positive number, **X**_(*j*,*ℓ*)_ and *y_j_* denote the voxel values of *ℓ* – *th* region and the class label of *j* – *th* snapshot, respectively. Further, **W**_*ℓ*_ = [**W**_(1,*ℓ*)_, **W**_(2,*ℓ*)_,…, **W**_(*j*,*ℓ*)_,…,**W**_(*τ*,*ℓ*)_ is the generated weights for predicting MVP model based on the *ℓ* – *th* active region. The classifier for *ℓ* – *th* region is also denoted by *η*_*ℓ*_, where all of these classifiers can be defined by *η* = {*η*_*L*1_,…,*η*_*ℓ*_,… *η*_*L*2_}. The final step in the proposed method is combining all classifiers (*η*) by Bagging [9] algorithm for generating the MVP final predictive model. Indeed, Bagging method uses the average of predicted results in (3.20) for generating the final result 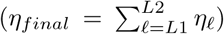 [9, 10]. Algorithm 3 shows the whole of procedure in the proposed method by using Leave-One-Out (LOO) cross-validation in the subject level.

##### Algorithm 3

The Proposed Method by using (LOO)

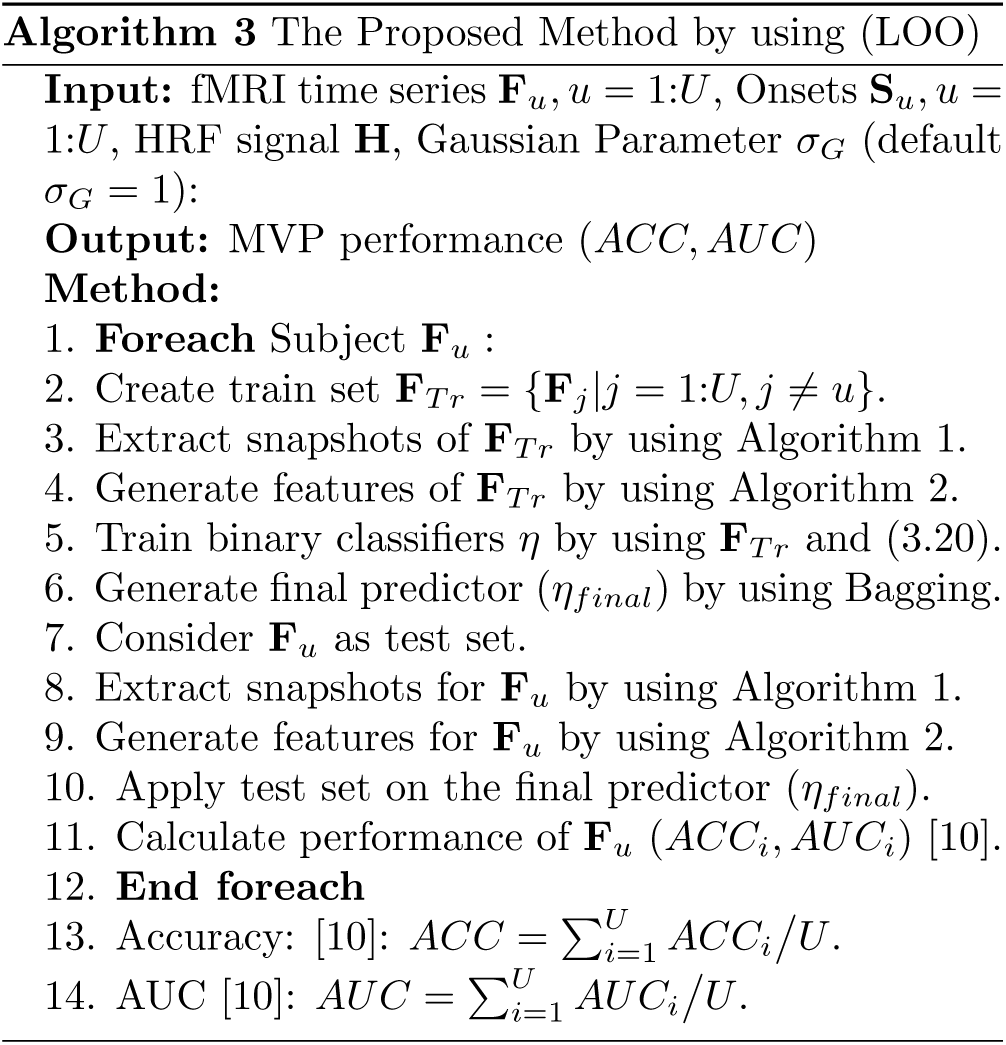

## 4 Experiments

### 4.1 Data Sets

This paper utilizes three data sets, shared by openfmri.org, for running empirical studies. As the first data set, ‘Visual Object Recognition’ (DS105) includes *U* = 71 subjects. It also contains *p* = 8 categories of visual stimuli, i.e. gray-scale images of faces, houses, cats, bottles, scissors, shoes, chairs, and scrambles (nonsense patterns). This data set is analyzed in high-level visual stimuli as the binary predictor, by considering all categories except nonsense photos (scramble) as objects. Please see [5, 6, 11, 14, 26] for more information. As the second data set, ‘Word and Object Processing’ (DS107) includes *U* = 98 subjects. It contains *p* = 4 categories of visual stimuli, i.e. words, objects, scrambles, consonants. Please see [27] for more information. As the last data set, ‘Multi-subject, multi-modal human neuroimaging dataset’ (DS117) includes MEG and fMRI images for *U* = 171 subjects. This paper just uses the fMRI images of this data set. It also contains *p* = 2 categories of visual stimuli, i.e. human faces, and scrambles. Please see [28] for more information. These data sets are separately preprocessed by SPM 12 (6685) (www.fil.ion.ucl.ac.uk/spm/), i.e. slice timing, realignment, normalization, smoothing. This paper employs the Montreal Neurological Institute (MNI) 152 T1 1mm as the reference image (**Ref**) in (3.11) for mapping the extracted snapshots to the standard space 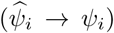. The size of this image in 3D scale is *X* = 182, *Y* = 218, *Z* = 182. Moreover, the *Talairach* Atlas [29] (including *L* = 1105 regions) in the standard space is used in (3.17) for extracting features. Further, all of algorithms are implemented in the MATLAB R2016b (9.1) on a PC with certain specifications^2^ by authors in order to generate experimental results.

### 4.2 Correlation Analysis

Figure 3 A, C, and E respectively demonstrate correlation matrix at the voxel level for the data sets DS105, DS107, and DS117. Further, Figure 3 B, D, and F respectively illustrate the correlation matrix in the feature level for the data sets DS105, DS107, and DS117. Since neural activities are sparse, high-dimensional and noisy in voxel level, it is so hard to discriminate between different categories in Figure 3 A, C, and E. By contrast, Figure 3 B, D, and F provide distinctive and informative representation, when the proposed method used the extracted features.

**Figure 3:**
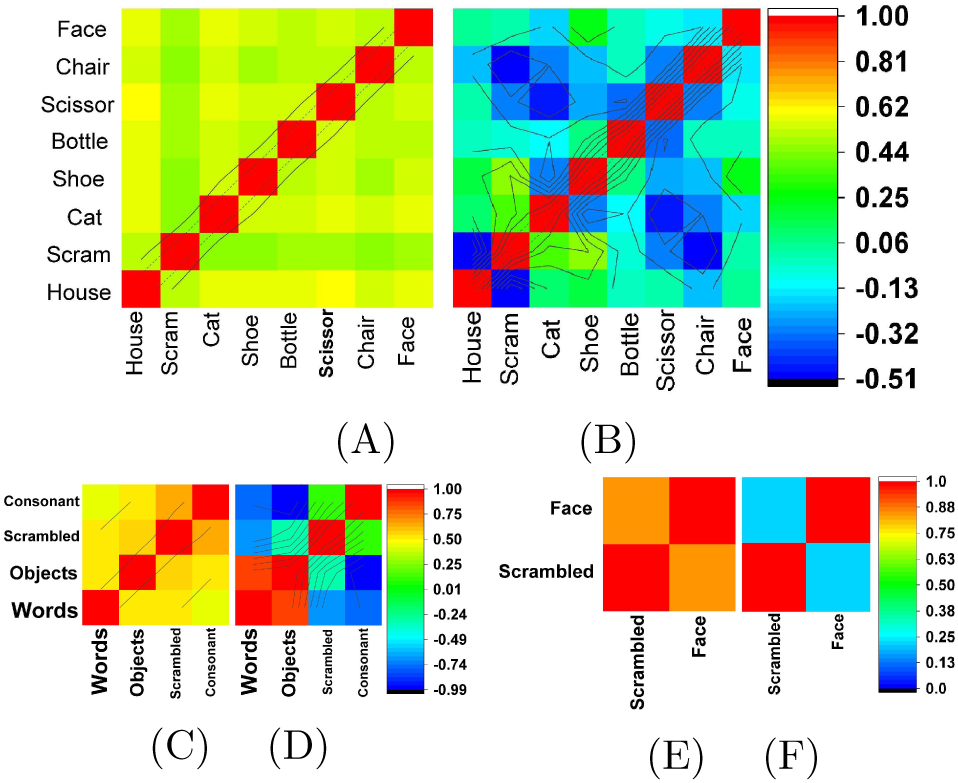
Correlation Matrix: for Visual Object Recognition (DS105) data set (A) in the voxel level, (B) feature level, for Word and Object Processing (DS107) data set (C) in the voxel level, (D) feature level, and for multi-subject, multi-modal human neuroimaging dat set (DS117) (E) in the voxel level, (F) feature le level.

### 4.3 Performance Analysis

The performance of our proposed method is compared with state-of-the-art algorithms, which were proposed for decoding the visual stimuli in the human brain. As a pioneer algorithm, our method is compared by SVM method [22], which is used in [6, 11] for decoding the visual stimuli. The performance of Graph Net [23] and Elastic Net [18] are reported as the most popular methods for fMRI analysis [3, 4, 16, 17, 19, 20]. Moreover, the performance of L1-Reg. SVM [8] is compared by the proposed method. The L1-Reg. SVM is recently employed by [3] as the most effective approach for decoding visual stimuli. Since this paper also applies L1-Reg. SVM for generating the predictive model in the level of ROIs, it can be considered as a baseline for comparing our feature space with the previous approaches. Lastly, Osher et al. [12] proposed a graph-based approach for creating predictors. Indeed, they employed the anatomical structure of the human brain for constructing graph networks. This paper compares the performance of the mentioned methods as well as the proposed method by using LOO cross-validation at the subject level. Further, the Gaussian parameter for smoothing the design matrix is considered *σ_G_* = 1. The effect of different values of this parameter on the performance of the proposed method will be discussed in the supplementary materials.

Table 1 and 2 respectively demonstrate the classification Accuracy and Area Under the ROC Curve (AUC) in percentage (%) for the binary predictors. These tables report the performance of binary predictors based on the category of the visual stimuli. All visual stimuli in the data set DS105 except nonsense photos (scramble) are considered as the object category for generating these experimental results. In addition, different categories of visual stimuli (including words, consonants, objects, and scrambles) in the DS107 are compared by using one-versus-all strategy. Moreover, face recognition based on neural activities is trained by using DS117 data set. Finally, all data sets are combined for generating predictive models for different categories of visual stimuli, i.e. faces, objects, and scrambles. As Table 1 and 2 demonstrate, the proposed algorithm has generated better performance in comparison with other methods because it provided a better representation of neural activities by exploiting the snapshots of the automatically detected active regions in the human brain. The last three rows in Table 1 and 2 illustrate the accuracy of the proposed method by combining all data sets. As depicted in these rows, the performances of other methods are significantly decreased. As mentioned before, it is the normalization problem. In addition, our framework employs the extracted features from the automatically detected snapshots instead of using all or a group of voxels, which can decrease noise and sparsity and remove high-dimensionality. Therefore, the proposed method can significantly decrease the time and space complexities and increase rapidly the performance and robustness of the predictive models.

**Table 1:** Accuracy of binary predictors

| Data Sets | SVM | Graph Net | Elastic Net | L1-Reg. SVM | Osher et al. | Proposed method |
| --- | --- | --- | --- | --- | --- | --- |
| DS105: Objects vs. Scrambles | 71.65±0.97 | 81.27±0.59 | 83.06±0.36 | 85.29±0.49 | 90.82±1.23 | **94.32±0.16** |
| DS107: Words vs. Others | 82.89±1.02 | 78.03±0.87 | 88.62±0.52 | 86.14±0.91 | 90.21±0.83 | **92.04±0.09** |
| DS107: Consonants vs. Others | 67.84±0.82 | 83.01±0.56 | 82.82±0.37 | 85.69±0.69 | 84.54±0.99 | **96.73±0.19** |
| DS107: Objects vs. Others | 73.32±1.67 | 77.93±0.29 | 84.22±0.44 | 83.32±0.41 | **95.62±0.83** | 93.07±0.27 |
| DS107: Scrambles vs. Others | 83.96±0.87 | 79.37±0.82 | 87.19±0.26 | 86.45±0.62 | 88.1±0.78 | **90.93±0.71** |
| DS117: Faces vs. Scrambles | 81.25±1.03 | 85.19±0.56 | 85.46±0.29 | 86.61±0.61 | **96.81±0.79** | 96.31±0.92 |
| ALL: Faces vs. Others | 66.27±1.61 | 68.37±1.31 | 75.91±0.74 | 80.23±0.72 | 84.99±0.71 | **89.99±0.31** |
| ALL: Objects vs. Others | 75.61±0.57 | 78.37±0.71 | 76.79±0.94 | 80.14±0.47 | 79.23±0.25 | **92.44±0.92** |
| ALL: Scrambles vs. Others | 81.92±0.71 | 81.08±1.23 | 84.18±0.42 | 88.23±0.81 | 90.5±0.73 | **95.39±0.18** |

**Table 2:** Area Under the ROC Curve (AUC) of binary predictors

| Data Sets | SVM | Graph Net | Elastic Net | L1-Reg. SVM | Osher et al. | Proposed method |
| --- | --- | --- | --- | --- | --- | --- |
| DS105: Objects vs. Scrambles | 68.37±1.01 | 70.32±0.92 | 82.22±0.42 | 80.91±0.21 | 88.54±0.71 | **93.25±0.92** |
| DS107: Words vs. Others | 80.76±0.91 | 77.91±1.03 | 86.35±0.39 | 84.23±0.57 | 87.61±0.62 | **91.86±0.17** |
| DS107: Consonants vs. Others | 63.84±1.45 | 81.21±0.33 | 80.63±0.61 | 84.41±0.92 | 81.54±0.31 | **94.03±0.37** |
| DS107: Objects vs. Others | 70.17±0.59 | 76.14±0.49 | 81.54±0.92 | 80.92±0.28 | **94.23±0.94** | 92.14±0.42 |
| DS107: Scrambles vs. Others | 80.73±0.92 | 77±1.01 | 85.79±0.42 | 83.14±0.47 | 82.23±0.38 | **87.05±0.37** |
| DS117: Faces vs. Scrambles | 79.36±0.33 | 83.71±0.81 | 83.21±1.23 | 82.29±0.91 | 94.08±0.84 | **94.61±0.71** |
| ALL: Faces vs. Others | 61.91±1.2 | 65.04±0.99 | 74.9±0.61 | 78.14±0.83 | 83.89±0.28 | **91.05±0.12** |
| ALL: Objects vs. Others | 74.19±0.92 | 77.88±0.82 | 73.59±0.95 | 79.45±0.77 | 75.61±0.89 | **89.24±0.69** |
| ALL: Scrambles vs. Others | 79.81±1.01 | 80±0.49 | 82.53±0.83 | 88.14±0.91 | 88.93±0.71 | **92.09±0.28** |

## 5 Conclusion

As a conjunction between neuroscience and computer science, Multivariate Pattern (MVP) is mostly used for analyzing task-based fMRI data set. There is a wide range of challenges in the MVP techniques, i.e. decreasing noise and sparsity, defining effective regions of interest (ROIs), visualizing results, and the cost of brain studies. In overcoming these challenges, this paper proposes Multi-Region Neural Representation as a novel feature space for decoding visual stimuli in the human brain. The proposed method is applied in three stages: firstly, snapshots of brain image (each snapshot represents neural activities for a unique stimulus) are selected by finding local maximums in the smoothed version of the design matrix. Then, features are generated in three steps, including normalizing to standard space, segmenting the snapshots in the form of automatically detected anatomical regions, and removing noise by Gaussian smoothing in the level of ROIs. Experimental studies on 4 visual categories (words, objects, consonants and nonsense photos) clearly show the superiority of our proposed method in comparison with state-of-the-art methods. In addition, the time complexity of the proposed method is naturally lower than the previous methods because it employs a snapshot of brain image for each stimulus rather than using the whole of time series. In future, we plan to apply the proposed method to different brain tasks such as risk, emotion and etc.

## Acknowledgment

We thank the anonymous reviewers for comments. This work was supported in part by the National Natural Science Foundation of China (61422204 and 61473149), Jiangsu Natural Science Foundation (BK20130034) and NUAA Fundamental Research Funds (NE2013105).

